# High dispersal levels and lake warming are emergent drivers of cyanobacterial community assembly over the Anthropocene in peri-Alpine lakes

**DOI:** 10.1101/419762

**Authors:** Marie-Eve Monchamp, Piet Spaak, Francesco Pomati

**Affiliations:** Eawag, Swiss Federal Institute of Aquatic Science and Technology, Department of Aquatic Ecology, 8600 Dübendorf, Switzerland; Swiss Federal Institute of Technology (ETH) Zürich, Institute of Integrative Biology, 8092 Zürich, Switzerland.

**Keywords:** Beta diversity, community phylogenetic structure, stochastic assembly, environmental forcing, meta-community, peri-Alpine lakes, eutrophication, climate change

## Abstract

Disentangling the relative importance of deterministic and stochastic processes in shaping natural communities is central to ecology. Studies on community assembly over broad temporal and spatial scales in aquatic microorganisms are scarce. Here, we used 16S rDNA sequence data from lake sediments to test for community assembly patterns in cyanobacterial phylogenies across ten European peri-Alpine lakes and over a century of eutrophication and climate warming. We studied phylogenetic similarity in cyanobacterial assemblages over spatial and temporal distance, and environmental gradients, comparing detected patterns with theoretical expectations from deterministic and stochastic processes. We found limited evidence for deviation of lake communities from a random assembly model and no significant effects of geographic distance on phylogenetic similarity, suggesting no dispersal limitation and high levels of stochastic assembly. We did not detect a significant effect of phosphorus and nitrogen levels on deviation of community phylogenies from random. We found however a significant decay of phylogenetic similarity for non-random communities over a gradient of air temperature and water column stability. We show how phylogenetic data from sedimentary archives can improve our understanding of microbial community assembly processes, and support previous evidence that climate warming has been the strongest environmental driver of cyanobacterial community assembly over the Anthropocene.

## Introduction

Understanding the mechanisms that determine changes in the structure and composition of natural communities over large spatial and temporal scales is critical, given the impacts that human activities have on biodiversity and ecosystem functions ^1^. The relative importance of stochastic and deterministic processes driving community assembly might vary over space and time: dispersal, demography, ecological interactions and evolutionary processes can all influence the structure of natural communities across scales ^2-7^. It is an on-going challenge to understand how anthropogenic environmental changes influence ecological and evolutionary mechanisms determining community assembly, particularly in aquatic microbes whose dispersal appears to have no boundaries ^8,9^.

Assembly studies focusing on ecological mechanisms in lake cyanobacterial communities have been scarce due to a lack of data at the appropriated spatial and temporal scale, despite the importance that these organisms have reached over the past decades for freshwater ecosystem functioning and services ^10^. Over the last century, the frequency and severity of cyanobacteria-dominated communities have increased in lakes and reservoirs worldwide despite remediation measures applied at the regional and international scale ^11,12^. Cyanobacterial blooms are often dominated by toxic species, and there is a global concern that environmental changes are promoting the geographic expansion of some potentially harmful taxa ^13,14^, due to a combined effect of increasing temperature and nutrient loads ^11,15,16^. Toxic species such as *Dolichospermum lemmermannii* and *Planktothrix rubescens* have indeed widened their geographic distribution, supporting the idea that some harmful cyanobacteria are spreading across temperate lakes ^16^. The role of geographic dispersal (where distance limits the establishment of new taxa) relative to turnover of taxa driven by environmental gradients has not been explicitly explored in the assembly of these globally important microorganisms.

In this study, we analysed cyanobacterial community composition data spanning over a hundred years and across ten lakes. We used 16S rDNA sequences from sediment cores of European peri-Alpine lakes (Fig. S1) that underwent directional environmental change characterised by climate warming and eutrophication ^16^. Our previous work has explored the patterns of long term change in alpha and beta diversity in lake cyanobacterial communities, showing a homogenization of assemblage composition at the regional scale ^16^. The aim of this study was to test for emergent deterministic (environment-driven) and stochastic (dispersal-driven) patterns in the phylogenetic structure of cyanobacterial assemblages across these different lakes of the same region ^2,17^.

We used a null-model that accounted for temporal changes in the size of the species pool to simulate random assembly. We then tested for deviation from random patterns as phylogenetic clustering and overdispersion: the tendency for taxa to co-occur with larger or smaller expectancy, respectively, than predicted by the null-model (Fig. 1A) ^18-22^. In most cases, dispersal-driven assembly would generate random taxa co-occurrence patterns, while environmental drivers would lead to deviation from random assembly ^19,21,23,24^. There can be interactions among assembly mechanisms that generate exceptions to these predictions ^25,26^. We however expect that comparison of phylogenetic structures to null-model simulations, combined with the patterns of community phylogenetic similarity across lakes and spatial or ecological distance, will allow us testing for deterministic and stochastic signatures in cyanobacterial community assembly.

**Figure 1.**
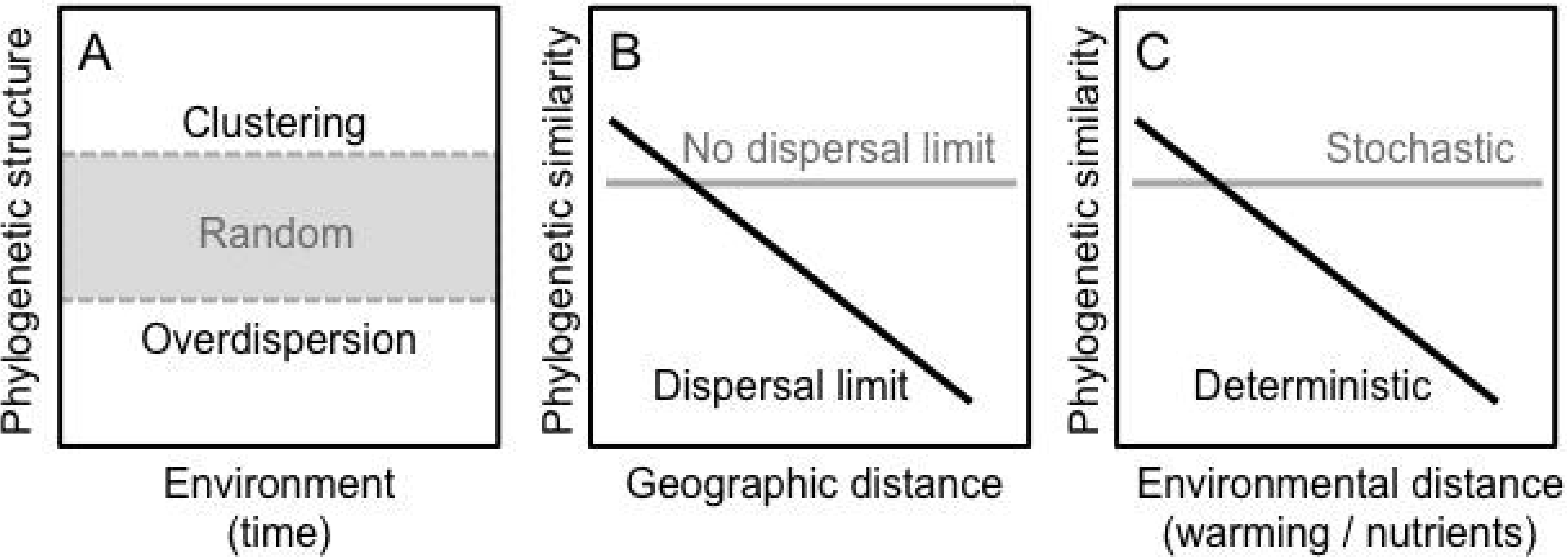
Schematic description of the theoretical expectations for phylogenetic community assembly within and between communities, based on stochastic (dispersal-driven) and deterministic (environment-driven) processes. **(A)** Taxa associations are analysed using their mean pairwise phylogenetic distance (MPD) for each date of each time-series and by comparing it to expected patterns from null-model simulations of random assembly (grey box): clustering and overdispersion (above and below the random expectation, respectively) signal communities that are composed of species phylogenetically closer or further apart than expected by chance, respectively, as a sign of deterministic processes. **(B)** Predicted patterns in phylogenetic community similarity depending on limitation (black line) and no limitation (grey line) in taxa dispersal among sites. **(C)** Predicted change in phylogenetic similarity for completely stochastic (grey) and deterministic (black) models of community assembly along an ecological environmental gradient (e.g. lake physics and chemistry).

Specifically, when dispersal of cyanobacterial taxa among lakes is not limited (Fig. 1B), we expect that phylogenetic community similarity will decrease over an environmental gradient, while no change is expected when the system is driven only by dispersal (Fig. 1C) ^2^. If there are barriers to dispersal of cyanobacteria, we predict differences in similarities among lake communities that are only dependent on the geographic distance (Fig. 1B), and no effects driven by ecological gradients (Fig. 1C) ^2^. The environmental-driven decrease in phylogenetic community similarity will not be influenced by dispersal limitation (Fig. 1B), and will vary deterministically as a consequence of the gradient itself (Fig. 1C) ^2^. This is because we expect that, under environment-driven assembly, the turnover of taxa along the ecological gradient will determine community structure in each lake. Here, we investigated whether cyanobacterial community phylogenetic structures within and across lakes over time matched these expectations from assembly processes, and what patterns dominate.

## Results

### Community phylogenetic structure

We calculated a standardised effect sizes (SES) of mean-pairwise distances (MPD) within each local community based on the comparison of the observed MPD values with the values a randomly assembled community (Methods). Based on the SES_MPD_ values, a large number of cyanobacterial communities did not show a phylogenetic structure that significantly differed from the null (random) expectation (Fig. 2). In fact, only two communities were significantly phylogenetically clustered, whereas eighteen out of seventy-six communities showed significant phylogenetic overdispersion, with SES_*MPD*_ values outside the 90% confidence interval of the null model simulation. Although the remaining forty-six communities analysed did not show significant signal of non-randomness, most (especially since the 1980s) of the SES_*MPD*_ values were negative, suggesting a tendency towards overdispersion. The samples that displayed significant non-random phylogenetic structure and their corresponding MPD and SES_*MPD*_ values are listed in Table S2 of the Supplementary Materials.

**Figure 2.**
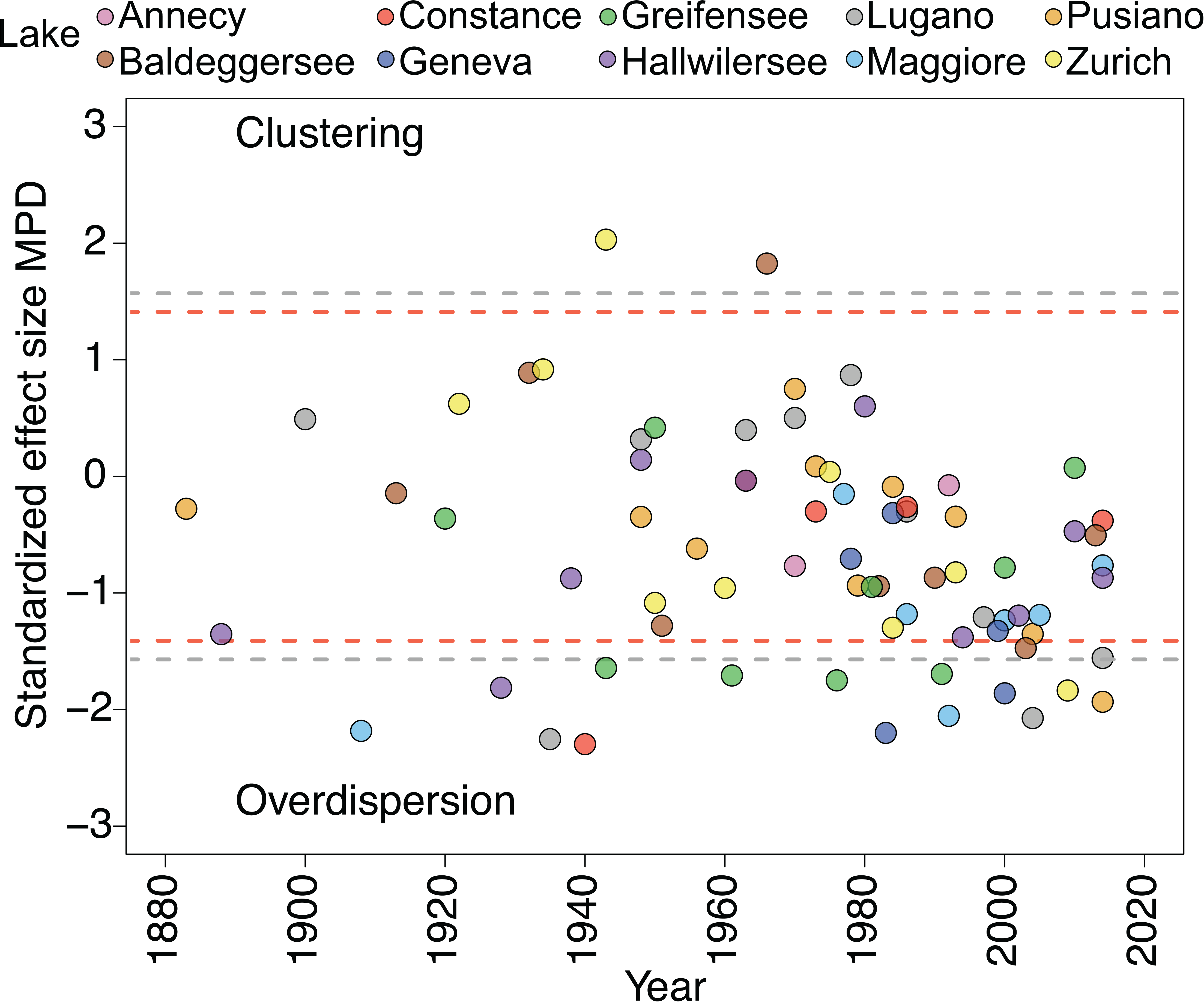
Time series of the standardised effect size of mean pairwise phylogenetic distances (SES_*MPD*_) values multiplied by -1 for the equivalent of the NRI index ^40^, calculated for each local community compared with null model simulations. The samples outside the central area delimited by the dashed lines show significant community structure (above or below the 95% [grey lines] and 90% [red lines] confidence intervals of the null-model simulation). Negative values signal phylogenetic overdispersion, whereas positive values signal phylogenetic clustering.

### Distance-decay relationships

We estimated the phylogenetic similarity (beta-diversity) across all pairs of communities reconstructed from the sedimentary archives of the ten lakes at each time-period and investigated the role of geographical and temporal distance (Fig. 3). Our analysis did not reveal an increase in phylogenetic beta-diversity with geographic distance (Fig. 3A), suggesting no dispersal limitation of cyanobacteria at the regional (peri-Alpine) scale. On the contrary, we observed for all lakes a clear decay of phylogenetic similarity along the temporal gradient representing the history of each lake (Fig. 3B), with the exception of lakes Geneva and Annecy (the latter due to insufficient data points) (Fig. S2).

**Figure 3.**
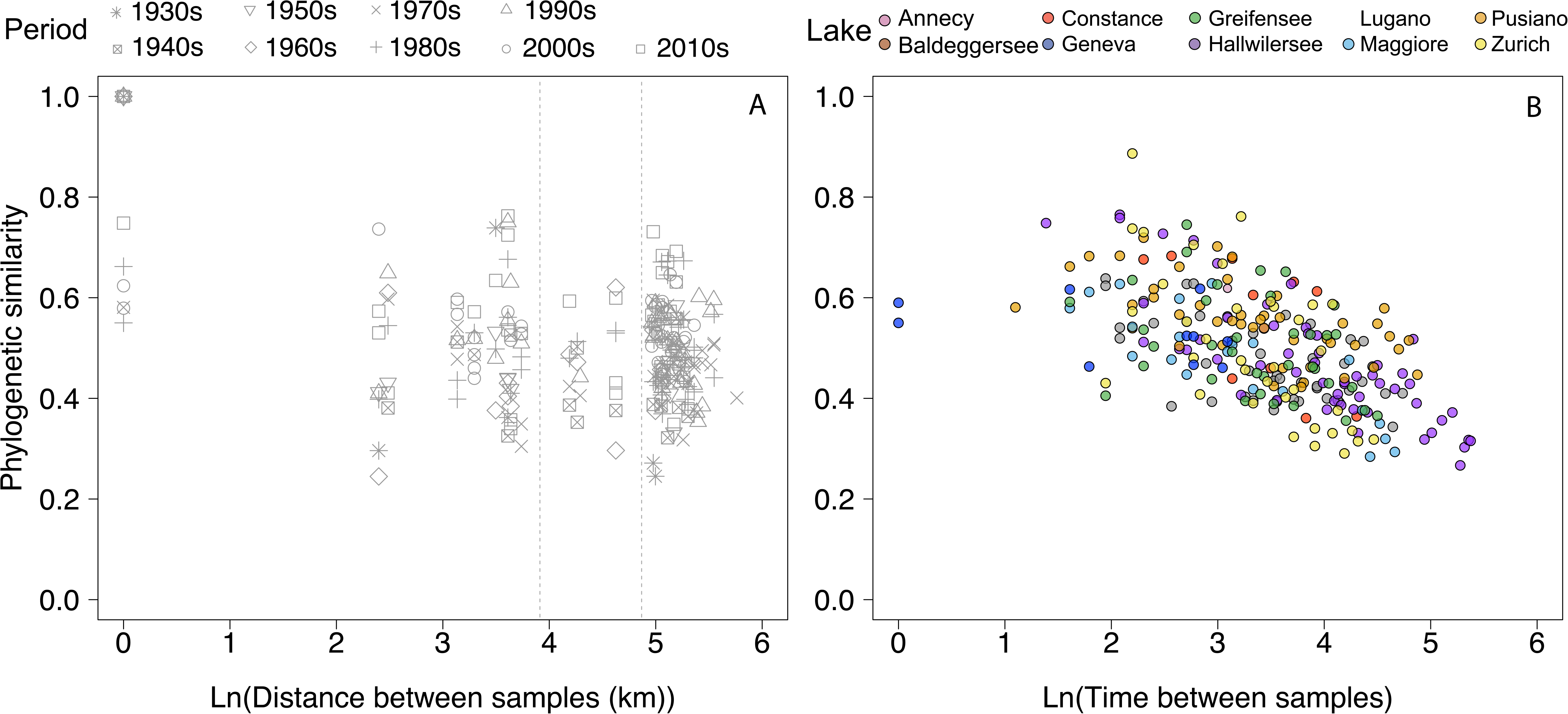
Distance-decay in cyanobacterial communities. **A)** Natural log-transformed phylogenetic (Unifrac) similarity between all pairwise cyanobacterial communities at each decade between the 1900s to the 2010s plotted against a gradient of natural log-transformed geographic distances between lakes. A null distance signifies that the pairwise phylogenetic dissimilarity was calculated between samples of the same lake at a given period. The vertical dashed lines mark the distances of 50 km and 130 km for reference (the absolute pairwise distances separating all lakes are summarized in Table S2 of Supplementary Information). **B)** Natural log-transformed Unifrac distances plotted against the natural log-transformed temporal gradient (years) for each lake (colour coding).

### Community similarity over environmental gradients

All non-random samples identified in Fig. 2 using the SES_*MPD*_ metric were used to investigate the role of chemical (TP and NO_3_^-^-N) and physical (air temperature and water column stability) drivers in explaining cyanobacterial community deviation from a random assembly. We found no evidence for a role of nutrients (TP and NO_3_^-^-N) in explaining non-random community structure (Fig. 4). On the other hand, OLS regression showed a significant decrease in community similarity along with both air temperature (*p* = 0.0003, adjusted R^2^ = 0.057) and SSI gradients (*p* = 0.019, adjusted R^2^ = 0.058) (Fig. 4). This suggests that communities in lakes characterized by similar physical characteristics related to lake water temperature are more similar in cyanobacterial community composition compared to lakes that display greater differences in temperature and stratification.

**Figure 4.**
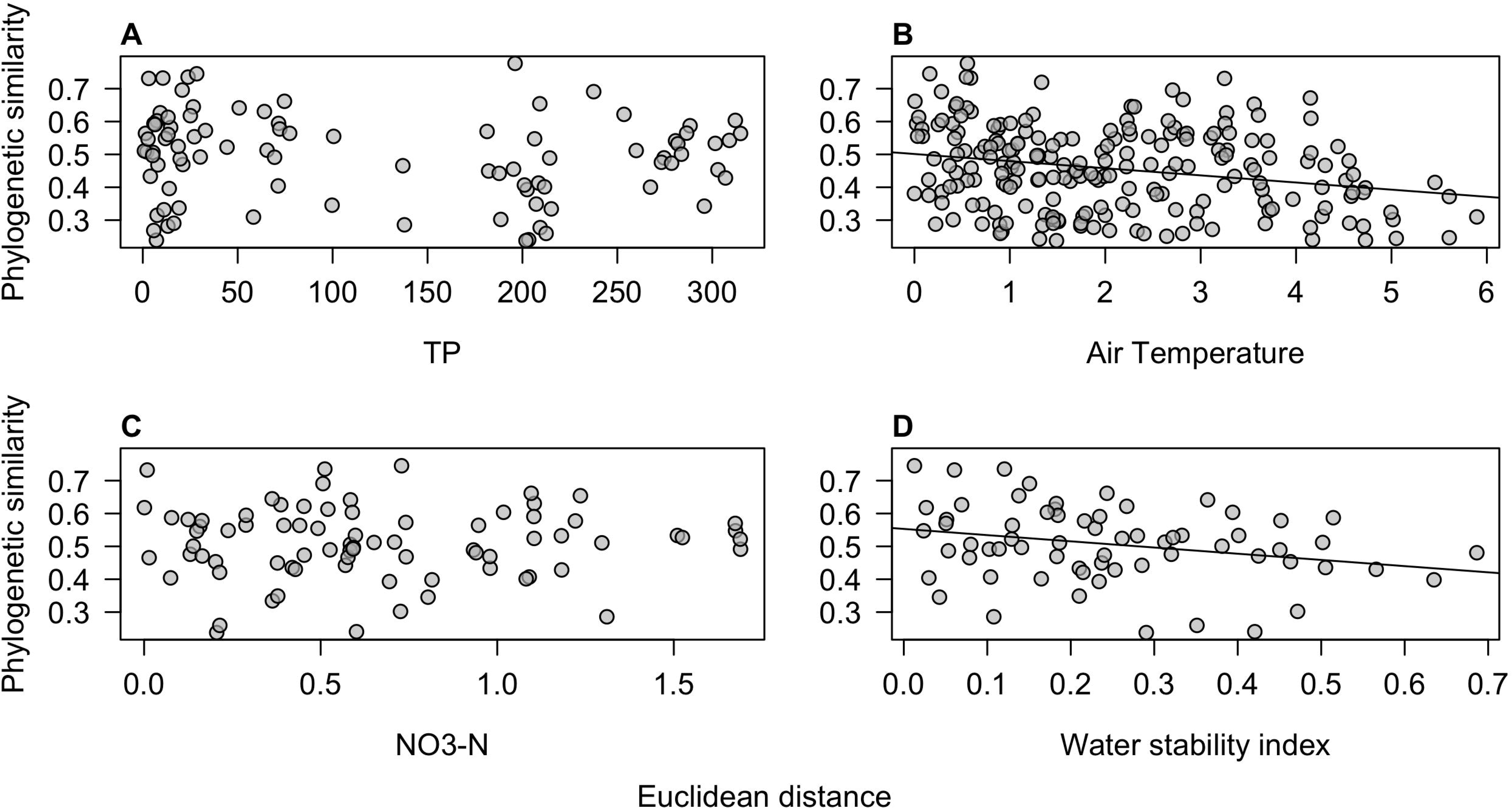
Cyanobacterial beta-diversity over environmental gradients. Phylogenetic (Unifrac) similarity between samples are plotted against environmental distances (euclidean) in **A)** TP (*n* = 104) and **B)** NO_3_ ^-^-N (*n* = 77) concentrations, **C)** air temperature (*n* = 209), and **D)** maximal annual Schmidt Stability Index (SSI) (*n* = 65). Only significant regressions (*p* ≤ 0.05) are shown. The number of samples in each analysis depends on the availability of lake monitoring data (details on the time-series of monitoring data are found in ^16^). Samples used in the regression analysis are the non-random communities (i.e., showing significant phylogenetic structure based on SES_*MPD*_ deviation from the null model expectation).

## Discussion

Most of the cyanobacterial communities obtained in this study from sedimentary archives did not show significant deviation from a random phylogenetic structure, suggesting a strong signal of stochastic (dispersal-driven) community assembly in lake cyanobacteria. Our previous work has shown that DNA-based reconstructions of cyanobacterial communities are robust ^16,27^, therefore the observed patterns are unlikely to be driven by biases in sedimentary DNA-based community reconstructions. On the contrary, it has been noted that a strong effect of stochastic relative to deterministic processes is expected in the community assembly of microalgae, particularly in highly productive (eutrophic) environments ^2^. The decay in phylogenetic similarity over time coupled with the lack of a geographic distance-decay relationship across lake communities (Fig. 3) suggest temporally dynamic communities with no limitation to dispersal at the regional (peri-Alpine) scale. In a recent study on genetic divergence among populations of a marine diatom, no significant relationship was found between genetic and geographic distances at regional and global scales ^9^. Most reports about microbial dispersal so far did no show clear evidence for geographic distance-decay patterns at the local (0 - 100 km) and regional (101 - 5,000 km) scales ^28^. The scale of distances in our study was not suited to capture dissimilarity changes among cyanobacterial communities along very large geographical distances (e.g. continental), where an effect of geographical isolation might emerge ^29^. Nevertheless, our research suggests that cyanobacterial communities present weak dispersal limitation among lakes of the same region, even around and across barriers such as the European Alpine mountain range.

Previous work has shown that communities of cyanobacteria have become more homogeneous in terms of composition across peri-Alpine lakes over the last decades, in favour of a few clades of bloom-forming and potentially toxic taxa ^14,16^. This could result in an increase of phylogenetic clustering over time, if the traits under selection by environmental changes are phylogenetically conserved ^19^. Our data show that the phylogenetic structure of assemblages is, in most cases, not clustered. Only two out of seventy-six tested communities displayed significant clustering, which strongly suggests that the environmental driving forces of cyanobacterial assembly in the lakes are not acting on phylogenetically conserved traits. Coloniality and buoyancy regulation are in fact multiphyletic traits that have been associated to the invading taxa, which belong to different cyanobacterial phylogenetic lineages within the orders *Chroococcales*, *Nostocales* and *Oscillatoriales* ^10,11,16^. The more prevalent signal of overdispersion in the community phylogenies might support this hypothesis of environmental selection for convergent traits.

Gradients of NO_3_^-^-N and TP concentrations across lakes did not significantly explain deviation from random assembly in the investigated cyanobacterial communities (Fig. 4). It is important to note that most of the lakes investigated here classify as meso-eutrophic to eutrophic ^30^ (see average P and NO_3_^-^-N concentrations reported by ^16^). Significant patterns of phylogenetic dissimilarity might emerge across communities characterized by a broader nutrient gradient, but this remains to be tested by surveys or experimentally. Rather, the difference in physical conditions, such as temperature and strength of the water column stratification, seemed in our study to explain a significant proportion of the observed variance in phylogenetic relatedness among lake cyanobacterial communities. The increasing trend in air temperatures, which has accelerated since the 1980s across the peri-Alpine region, has caused modifications in thermal characteristics of lakes, e.g., via changes in the duration and strength of the water column stratification ^16,31^, which in turn affects recirculation and availability of nutrients for phytoplankton growth ^10,22,32^. This effect has been amply documented and has favoured, as mentioned above, buoyant cyanobacterial forms that are able to control their vertical position in the water column to reach optimal nutrient and light conditions ^14,16,22,31,32^. Our findings therefore support previous evidence and suggest that climate warming is the strongest environmental driver of the assembly of lake cyanobacterial communities ^33^, and might select for specific traits such as, for example, buoyancy regulation.

In conclusion, this is the first study to our knowledge that explicitly tests for deterministic and stochastic assembly patterns in cyanobacterial communities across regional scales and over the Anthropocene. We successfully reconstructed the phylogenetic diversity in seventy-six cyanobacterial communities over a period of ca. 100 years using DNA from sediment cores collected in ten lakes. Our study shows that both stochastic (dispersal-driven) and deterministic (environmental-driven) processes are important in assembling cyanobacterial communities across lakes of the European peri-Alpine region. Cultural eutrophication and climate change are the most notable environmental factors favouring cyanobacterial growth, but the deterministic processes governing community assembly appeared in our study to be more significantly driven by lake warming. Our results confirm previous evidence ^16,33^ and expand our understanding of cyanobacterial community assembly processes. Knowledge about the relative importance of (potentially controllable) environmental drivers and (likely uncontrollable) dispersal of organisms in shaping the structure of cyanobacterial assemblages is important for the management of aquatic ecosystems whose services are threatened by an increasing prevalence of potentially toxic taxa.

## Materials and methods

### Data collection

We used high-resolution 16S rDNA sequence data from ten European peri-Alpine lakes, spanning between early 1900s and 2016, to estimate phylogenetic diversity of cyanobacterial communities ^16^. Briefly, sediment cores were collected in ten lakes between 2013 and 2016 using a gravity corer, and layers were dated by varve counting and, in most cases, with radionuclide (Pb^210^, Cs^237^) measurements ^16,27^. Based on the sediment age models, sediment sub-samples were collected at various depths in cores from each lake to capture the cyanobacterial community composition over the last ∼100 years. DNA was extracted from bulk sediments in a clean laboratory facility, and the DNA extract were used for PCR and high-throughput sequencing of the cyanobacterial 16S rRNA gene (Table S1) on a MiSeq Illumina platform as previously described ^16,27^.

The clean, primmer-trimmed sequences were clustered in operational taxonomic units (OTUs) with a 97% threshold of sequence similarity in QIIME ^34^ using the UPARSE workflow ^35^. PyNast ^34^ and the Greengenes microbial sequence database ^36^ were used for sequence alignment, and FastTree ^37^ was used to estimate a phylogeny based on maximum-likelihood containing all OTUs found in the lakes. OTUs were taxonomically assigned with a confidence threshold of 85% and the ones assigned to phyla other than photosynthetic cyanobacteria were removed from the dataset. Each sample was rarefied to 2,744 sequences (cyanobacteria only) prior to phylogenetic analyses.

The physical (air temperature) and chemical (nitrogen [NO_3_^-^-N], total phosphorus [TP]) data were collected over several decades consist of several decades of monitoring of the ten lakes ^16^ and the annual maximal Schmidt Stability Index (SSI; the maximal strength of the water column stratification) was derived from water temperature and hypsometry data ^16,38^. Euclidean distances for each environmental variable were calculated among lakes to derive environmental gradients used in the linear models in the R package ‘vegan’ version 2.4.4 ^39^.

### Phylogenetic analyses

To derive the phylogenetic structure of each community, we quantified the mean pairwise phylogenetic distance (MPD) ^40^ using *mpd* and *ses.mpd* in the package ‘picante’ version 1.6.2 for R ^41^ and used null-model simulations of random assembly that account for temporal changes in the size of the species pool ^21^. We calculated a standardised effect size of MPD (SES_*MPD*_) within each local community subtree based on the comparison of the observed MPD values with the values in the random distribution using 999 randomisations of the species at the tip of the phylogenetic tree, while species richness was maintained ^41-43^: SES_*MPD*_ = Mean(MPD_*Observed*_ -MPD_*Randomized*_)/SD(MPD_*Randomized*_). The SES_*MPD*_ values were multiplied by −1 to be equivalent to the net relatedness index (NRI) ^40^.

We derived the Unifrac phylogenetic distance ^44^ between all pairs of samples using the *dist* function in the Bioconductor package ‘phyloseq’ ^45^ based on the OTU table and the fasta files from amplicon sequencing. The geographic distance-decay relationship was measured on binned communities each representing a period of one decade (from the 1930s to the 2010s; decades 1900s, 1910s and 1920s were excluded due to insufficient number of samples). The binning was done to remove the factor time from the analysis, as it would introduce a bias when comparing multiple samples from single lakes over time. The temporal distance-decay pattern of phylogenetic similarity was studied by plotting log-transformed Unifrac distances against natural log-transformed time distances (years) for each lake in the dataset. Significance of the distance-decay relationship at each decade was tested using Mantel tests in the R package ‘ade4’ with a significance threshold of *p* ≤ 0.05. To test whether the environment (physical and chemical parameters) play a role in driving community assembly, we used the samples that were identified as phylogenetically non-random (i.e., those which SES_*MPD*_ values were outside the 90% confidence interval of the null model simulation) in linear ordinary least square (OLS) regressions where physical and chemical lake data were the explanatory variables.

## Acknowledgments

This work was supported by the Swiss Enlargement Contribution, project IZERZ0 – 142165 ’CyanoArchive’ to P.S. in the framework of the Romanian–Swiss Research Programme.

## Authors contributions

M-E.M., F.P. and P.S. designed the study. M-E.M. collected data and performed data analysis. M-E.M. and F.P. wrote the manuscript, which was commented and edited by P.S.

## Conflict of Interest

The authors declare no competing financial interests.

